# High cooperativity in negative feedback can amplify noisy gene expression

**DOI:** 10.1101/125914

**Authors:** Pavol Bokes, Yen Ting Lin, Abhyudai Singh

## Abstract

Burst-like synthesis of protein is a significant source of cell-to-cell variability in protein levels. Negative feedback is a common example of a regulatory mechanism by which such stochasticity can be controlled. Here we consider a specific kind of negative feedback, which makes bursts smaller in the excess of protein. Increasing the strength of the feedback may lead to dramatically different outcomes depending on a key parameter, the noise load, which is defined as the squared coefficient of variation the protein exhibits in the absence of feedback. Combining stochastic simulation with asymptotic analysis, we identify a critical value of noise load: for noise loads smaller than critical, the coefficient of variation remains bounded with increasing feedback strength; contrastingly, if the noise load is larger than critical, the coefficient of variation diverges to infinity in the limit of ever greater feedback strengths. Interestingly, high-cooperativity feedbacks have lower critical noise loads, implying that low-cooperativity feedbacks in burst size can be preferable for noisy proteins. Finally, we discuss our findings in the context of previous results on the impact of negative feedback in burst size and burst frequency on gene-expression noise.

## 1 Introduction

Due to the low molecular copy numbers involved, gene expression at a single cell level exhibits a considerable degree of noisiness (Elowitz et al, 2002; Blake et al, 2003). Noise in gene expression is transmitted downstream to the final product of gene expression, the protein, whose concentration can fluctuate widely in time (Golding et al, 2005; Suter et al, 2011). The fluctuations are particularly pronounced if the protein is produced in bursts of multiple molecule copies at a single time (Ozbudak et al, 2002; Dar et al, 2012). The temporal fluctuation of protein concentration manifests itself in a population of isogenic cells as a cell-to-cell variability (Taniguchi et al, 2010). Mathematical modelling can be used to analyse the impact of mechanisms involved in gene expression on the population distribution of protein concentration (Shahrezaei and Swain, 2008).

Many proteins control their own expression with negative feedback (Rosenfeld et al, 2002). By reducing the protein synthesis rate when the protein is in excess, negative feedback reduces the mean protein level and can also decrease the protein noise (Thattai and van Oudenaarden, 2001). Nevertheless, in the case of a strong negative feedback, any decrease in noise due to improved regression to the mean can be eclipsed by an increase due to low reaction frequencies and/or low copy numbers (Singh and Hespanha, 2009). Additionally, the noise-reducing effect of time-averaging of the downstream elements, such as the promoter state or the mRNA copy number, can be cancelled under strong feedback, thus enhancing bursting dynamics and increasing the protein noise even further (Stekel and Jenkins, 2008; Grönlund et al, 2013). Bursting in itself imposes limitations on the noise-reducing ability of a strong negative feedback: individual bursts can grow too large to remain under control of a strong feedback, which is sensitive to small changes in protein copy numbers (Scott et al, 2007; Wang et al, 2014; Bokes and Singh, 2016). It is this final mechanism — the loss of control over large bursts — that we shall investigate in this paper using a minimalistic model which purposely disregards other mechanisms.

Feedback in gene expression can operate on the transcriptional or post-transcriptional level. In transcriptional feedback, the protein regulates the rate with which its mRNA transcript is synthesised (Becskei and Serrano, 2000). Provided that the mRNA life span is short and the translation of mRNA is fast, the transcription of an mRNA molecule is followed by a quick burst of protein translation (Cai et al, 2006). Transcriptional feedback thus regulates the rate, or frequency, with which such translational bursts occur (Friedman et al, 2006). Post-transcriptional feedback, on the other hand, regulates the size of translational bursts (Schikora-Tamarit et al, 2016). As a specific example, RNA binding proteins reduce the size of translational bursts by enhancing the degradation of their mRNA transcript (Yates and Nomura, 1981; Swain, 2004; Schikora-Tamarit et al, 2016). Previous studies indicate that post-transcriptional feedback outperforms transcriptional feedback in terms of its noise-reducing capability (Swain, 2004; Singh, 2011; Bokes and Singh, 2016).

Production delay can lead to dramatic changes in qualitative behaviour of feedback systems (Mackey et al, 1977). Deterministic systems can develop sustained oscillations via a supercritical Hopf bifurcation from a destabilised steady state once a threshold value of the delay is exceeded (Murray, 2003). Stochastic systems with negative feedback become more variable upon the inclusion of a small delay (Lafuerza and Toral, 2011) and exhibit increasingly regular oscillations prior to the onset of the Hopf bifurcation (Barrio et al, 2006). Gene-expression feedback is necessarily delayed, since both the transcription of a gene into an mRNA and the translation of an mRNA into a protein involve a series of multistep processes before they are completed (Alberts et al, 2002). Large delays in gene expression of a negatively regulating transcription factor can lead to physiologically consequential sustained oscillations (Monk, 2003). In this paper we shall see that the ability of strong negative feedback in burst size to contain gene-expression noise is compromised by an infinitesimally small delay that is implicitly included in our model.

While some gene-expression models are exactly solvable (Biancalani and Assaf, 2015; Smith and Shahrezaei, 2015; Kumar et al, 2014; Grima et al, 2012; Bokes et al, 2012; Ramos et al, 2011; Jia and Kulkarni, 2010; Peccoud and Ycart, 1995), one has often to resort to asymptotic methods to obtain tractable results (Dessalles et al, 2017; Popovic et al, 2016; Be’er and Assaf, 2016; Newby, 2015; Bruna et al, 2014; Leier et al, 2014; Platini et al, 2011). Small-noise approximations, which linearise the model around a deterministic steady state, play an important role among the available asymptotic approaches (Elf and Ehrenberg, 2003; Paulsson, 2004; Komorowski et al, 2013; Wang et al, 2014; Thomas and Grima, 2015; Cardelli et al, 2016). However, expressions obtained by a small-noise approximation need not be uniformly valid across the whole parameter space of the model. Bokes and Singh (2016) used an alternative approximation in the regime of strong negative feedback to gain a thorough understanding of the mean and noise behaviour for an exactly solvable model across its parameter space. Here we shall apply small-noise and strong-feedback approximations in a model for which an exact distribution is, to our best knowledge, unavailable.

We adapt a popular framework which models the protein concentration by a continuous-state drift-jump Markov process in which the deterministic drift represents protein decay and the stochastic jumps model the synthesis of protein in bursts (Friedman et al, 2006; Bokes et al, 2013; Lin and Doering, 2016). In the original formulation, the decay of protein is exponential, the jumps occur randomly in time with a given intensity, and their sizes are drawn at random from an exponential distribution (Friedman et al, 2006; Bokes et al, 2013; Lin and Doering, 2016). Extensions to the original formulation also consider non-exponential decay (Mackey et al, 2013; Bokes and Singh, 2015; Soltani et al, 2015), non-exponential bursting distributions (Jedrak and Ochab-Marcinek, 2016b), multiple gene copies (Jedrak and Ochab-Marcinek, 2016a), and multiple-component systems (Yvinec et al, 2014; Mackey and Tyran-Kaminska, 2015; Lin and Galla, 2016; Pájaro et al, 2017). Most previous studies include feedback in burst frequency: the jump intensity depends on the current level of protein. Bokes and Singh (2016) introduced a particular form of feedback in burst size, in which the hazard rate for burst termination depends on the current protein concentration. Here we shall consider a different formulation of feedback in burst size, in which the protein concentration immediately before a jump determines the mean jump size. We shall argue that the two formulations of burst-size feedback differ in the inclusion or omission of an infinitesimally small delay in protein synthesis. As we shall see, the two formulations have strikingly different noise-reduction capabilities.

The modelling framework is first introduced in Section 2 for the simple case of a constitutively expressed protein and then extended by feedback in burst-size in Section 3. The fundamental features of the model behaviour are illustrated in Section 4 using stochastic simulations. Sections 5 and 6 contain a systematic asymptotic analysis of the model behaviour in the small-noise and strong-feedback regimes. Section 7 compares the current model to its alternatives in (Bokes and Singh, 2016) and Section 8 concludes with a summary and discussion of the main findings.

## 2 Constitutive case

We model the dynamics of protein concentration *x*(*t*) by a Markovian drift-jump process (Schuss, 2009), whose trajectories look like the one in Figure 1, left panel. Except for a countable number of discontinuities, the process *x*(*t*) decays deterministically with a rate constant *γ*. The jumps (bursts of protein expression) occur randomly in time with a constant burst frequency *a*; the waiting time between two consecutive bursts is exponentially distributed with mean 1/*a*. The size of the discontinuity (burst size) is always positive and randomly chosen from an exponential distribution with mean *b* (Friedman et al, 2006).

**Figure 1:**
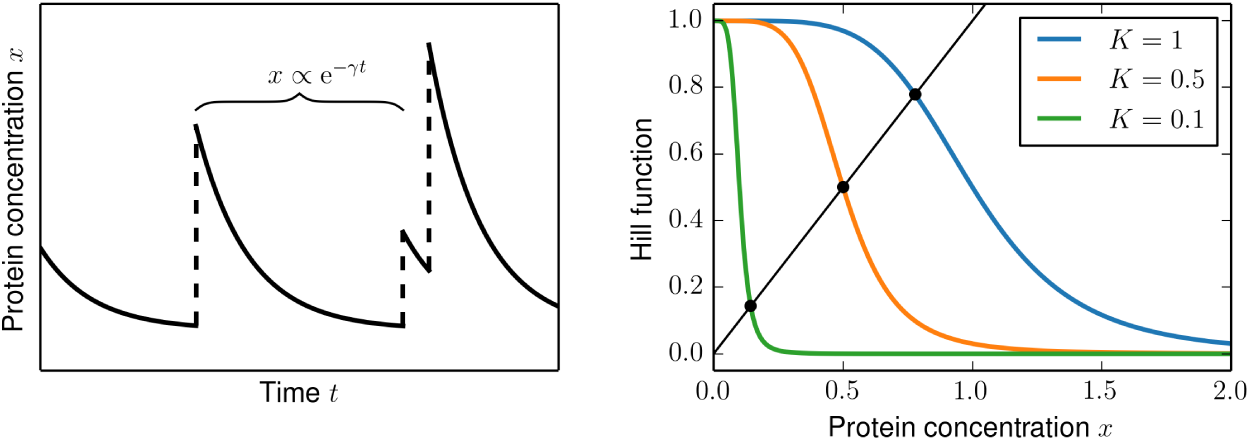
*Left:* A sample trajectory of the drift-jump process modelling the dynamics of protein concentration. *Right:* Hill functions (1 + (*x*/*K*)^5^)^−1^ and their fixed points for selected values of *K*.

There are three dimensional parameters in the model so far: decay rate constant *γ* (units of time^−1^), burst frequency *a* (units of time^−1^), and mean burst size *b* (units of concentration). We choose the units of time and concentration so that these parameters take the numerical values of

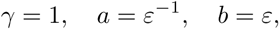
where *ε* is a positive dimensionless real number (Fig. 2).

**Figure 2:**
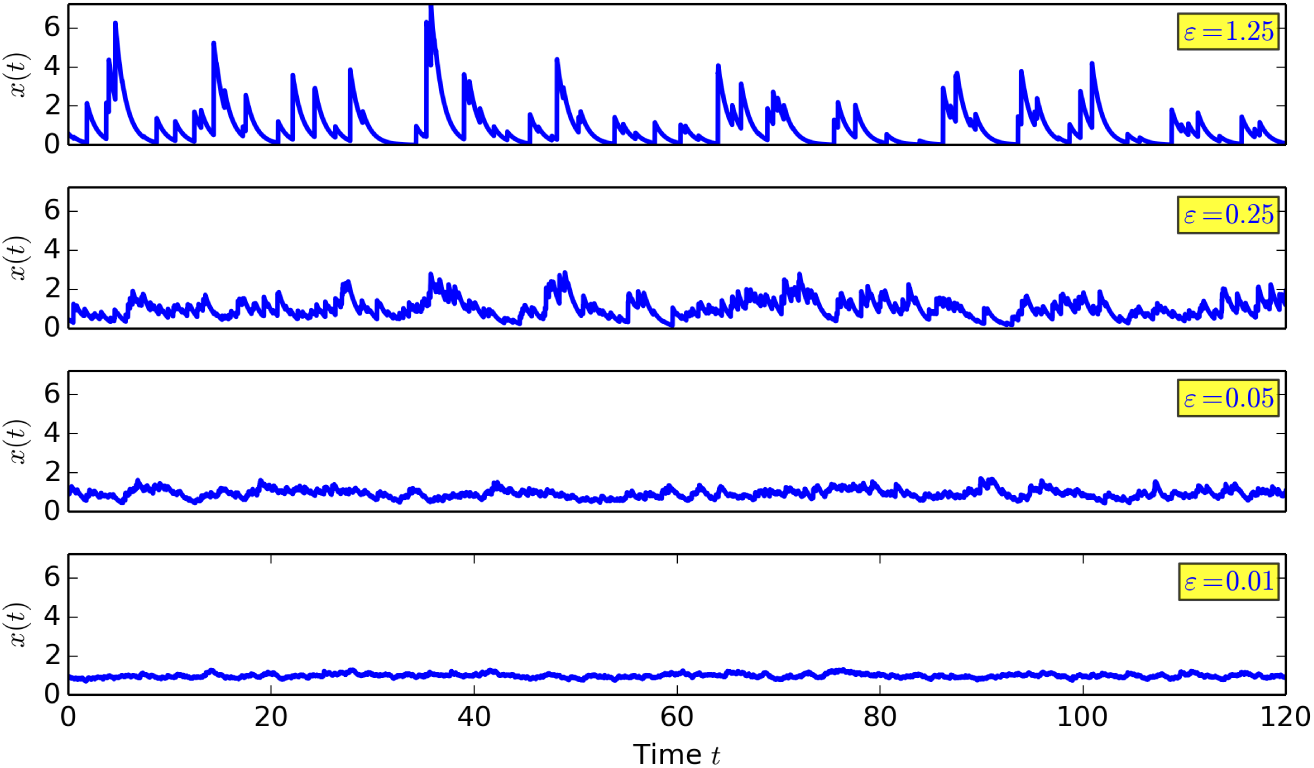
Sample trajectories for a constitutively expressed protein for selected values of *ε*.

The probability density function (pdf) *p*(*x, t*) of the Markov process *x*(*t*) satisfies a master equation (cf. Schuss, 2009)

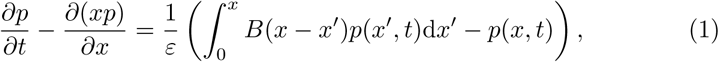
where

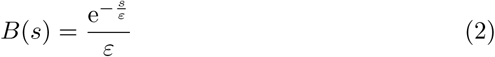
is the exponential burst-size probability density function. The normalised stationary solution to (1)–(2) is given by the gamma probability distribution (cf.Friedman et al, 2006)

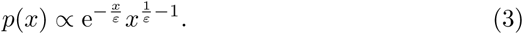

Here and elsewhere we shall use the same symbol for the independent variable of a pdf and a random variable (or a stationary process) associated with the pdf. For the gamma protein distribution in (3), the mean and variance are given by

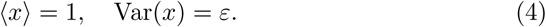

We henceforth call *ε* the *noise load* and say that the model operates in the *small-noise regime* if *ε* ≪1. We can normalise the protein concentration by writing

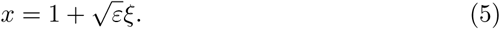

The stationary pdf satisfies

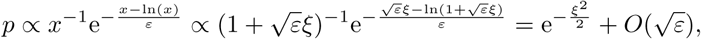
implying that in the small-noise regime the protein concentration is approximately normally distributed.

## 3 Feedback in burst size

We incorporate negative autoregulation of burst size into the model of Section 2 by making the mean burst size *b* depend on the concentration *x* of the protein at the time *immediately before* the burst occurs. Specifically, we shall consider a Hill-type dependence

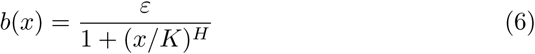
of the mean burst size on the protein concentration (Fig. 1, right panel). The critical concentration *K*, which gives the amount of protein that is required to halve the expected burst size, is a reciprocal measure of feedback strength; we say that the model operates in the *strong-feedback regime* if *K* ≪ 1. The cooperativity coefficient *H* determines how steeply the feedback responds to changes in protein concentration.

The master equation for the pdf *p*(*x*,*t*) of the process *x*(*t*) with negative feedback in burst size is given by

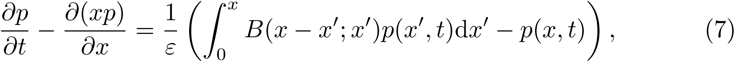
where

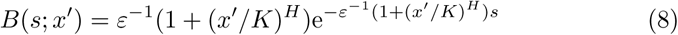
gives the burst-size density on condition of *x′* protein being present immediately before the burst. Notably, the convolution on the right-hand side of (1) of the pdf with an exponential density (2) is replaced in (7) by a more challenging integral kernel (8). Being unaware of the availability or otherwise of explicit solutions to (7)–(8), we shall resort to asymptotic methods to analyse its normalised stationary solution in the small-noise (*ε* ≪ 1) and strong-feedback (*K* ≪ 1) regimes.

While solving the master equation becomes problematic after the inclusion of feedback in burst size, simulating stochastic trajectories remains extraordinarily simple. Algorithm 1 loops over a basic procedure which consists of drawing the waiting time *τ* until the next burst (Line 4), setting the trajectory to an exponential in between the bursts (Line 5), and drawing the next burst’s size (Line 6). The waiting time and burst size are both exponentially distributed and can be drawn using the inversion sampling method by multiplying their respective mean values by –ln*u*, where *u* is drawn independently of any other drawings from the uniform distribution in the unit interval.

### Algorithm 1: Simulation algorithm

~~~
**Require:** Parameter values (*ε, H, K*).
**Return:** A sample trajectory *x*(*t*), *t* ≥ 0.
 1: Initialise the current time and state: *t* ← 0; *x*(0) ← 0.
 2: **loop**
 3:   Draw *u* and *ũ* from the standardised uniform distribution.
 4:   Set *τ* ← –*ε*ln*u*.
 5:   Set *x*(*t* + *s*) ← *x*(*t*)*e^−s^* for 0 < *s* < *τ*.
 6:   Set *x*(*t* + *τ*) ← *x*(*t*)*e^−τ^* 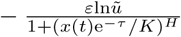.
 7:   Update the current time *t*  ← *t* + *τ*.
 8: **end loop**
~~~

## 4 Critical noise load

Stochastic simulations by Algorithm 1 allow us to investigate the response of protein noise to the strengthening of feedback in burst size. We choose to quantify protein noise by the (squared) coefficient of variation (CV^2^), which is defined as the ratio of variance over the mean squared, i.e.

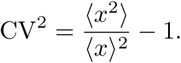

By (4), the squared coefficient of variation reduces to the value of the noise load *ε* in the absence of feedback (*K* → ∞). The ratio of the CV^2^ of a regulated protein to *ε*,which we shall refer to as the relative CV^2^, gives the fold increase in protein noise following the introduction of feedback.

Depending on whether the noise load *ε* is less or greater than a critical value *ε_c_* = 1/2*H*, we observe two different responses of protein noise to strengthening feedback. If the noise load is subcritical (*ε* < *ε_c_*), the noise in protein concentration remains bounded however strong the feedback may be (Fig. 3); for supercritical noise loads (*ε* > *ε_c_*), the noise diverges as *K* tends to zero (Fig. 4). The vertical (concentration) axes in the panels of Figures 3 and 4 extend from zero to a constant multiple (set to 2 in Fig. 3 and to 10 in Fig. 4) of the protein mean; the apparent variability in the sample trajectory then directly corresponds to the numerical value of the coefficient of variation, which we report in the inset of each panel. Each coefficient of variation is calculated from a single run of simulation which is much longer than the short stretch shown in the individual panels (see Appendix A for details).

**Figure 3:**
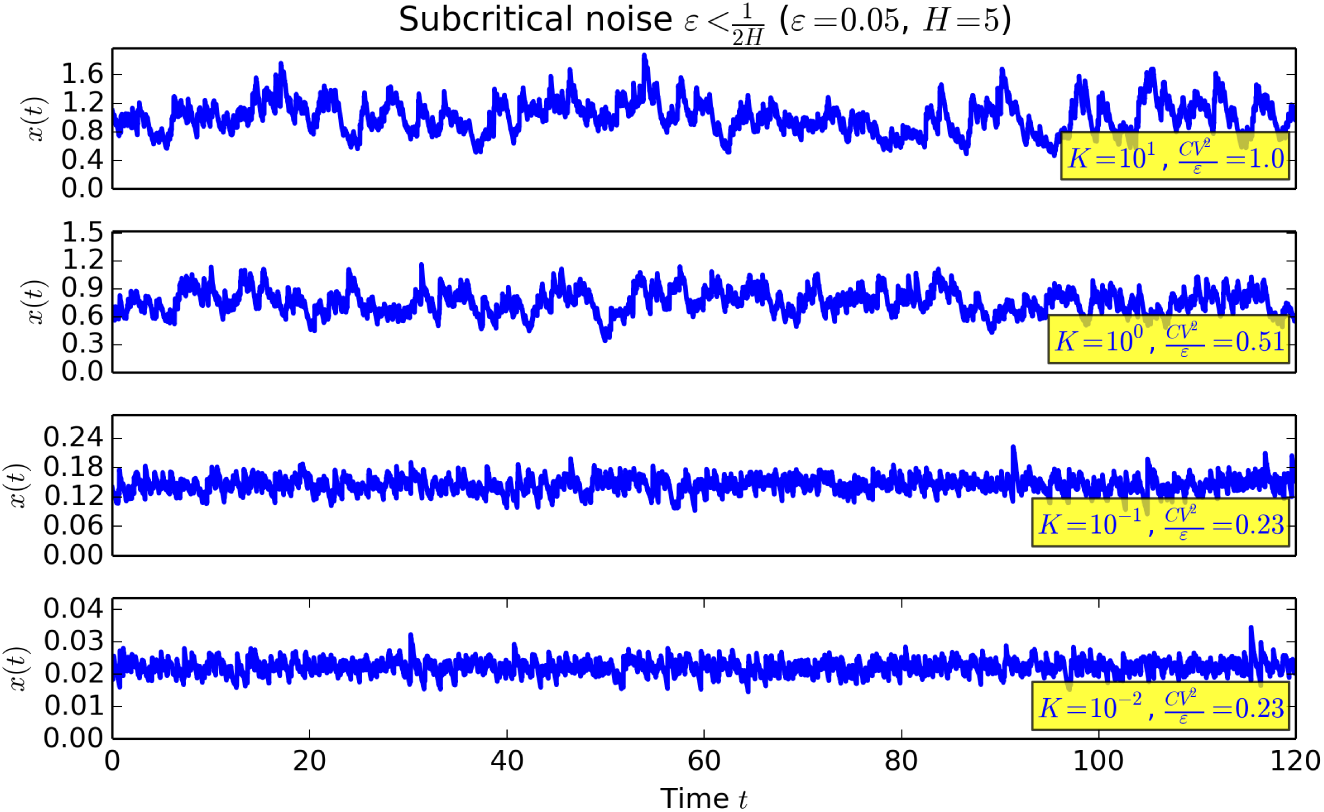
For subcritical noise loads, protein noise remains bounded even under stringent feedback conditions.

**Figure 4:**
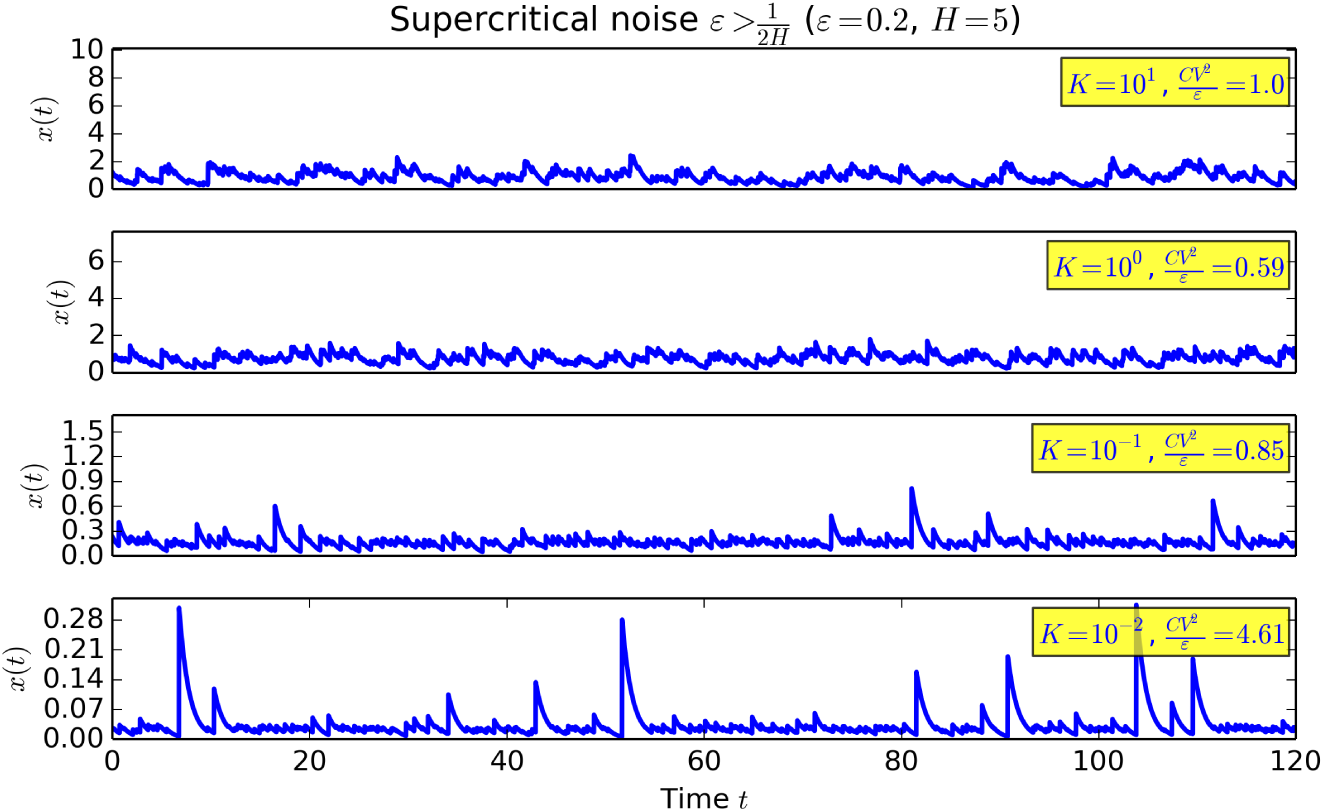
For supercritical noise loads, protein noise diverges for increasingly strong feedback.

As we shall demonstrate in Section 6 systematically using asymptotic analysis, the protein mean 〈*x*〉, the mean square 〈*x*^2^〉, and the variance all tend to zero, in subcritical and supercritical cases alike, as feedback strengthens. The increase of the coefficient of variation occurs because in supercritical cases the squared mean 〈*x*〉^2^ converges faster to zero than the mean square 〈*x^2^*〉.

Steeper (more cooperative) feedbacks have lower critical noise loads. If we take the cooperativity coefficient *H* to infinity, the critical noise load becomes zero: any positive noise load is supercritical. The Hill function (6) formally reduces for *H* = ∞ to a step function which is equal to *ε* for 0 < *x* < *K* and is zero for *x* > *K*. Under such step-wise regulation, non-zero bursts occur only if *x* is smaller than *K* (i.e. very small), and their size is independent of *K* (i.e. large); after one such large burst no further bursts can occur—or we may say that they still do but their size is necessarily—zero until the protein concentration slowly decays back beneath *K*. Clearly, however small, but fixed, *ε* may be, these infrequent but large bursts will drive the variability to infinity if *K* is lowered sufficiently. However, for finite cooperativities a non-zero burst would typically fire well before the concentration decays down to levels of O(*K*), and would be of a lesser size as a result: it is exactly this mechanism that enables low-cooperativity feedbacks to cope with ever increasing feedback strengths; the underlying scaling will be identified and analysed by matched asymptotics in Section 6. Prior to doing that, we perform in the following Section 5 a traditional small-noise analysis, obtaining explicit mean and noise results, which are valid as *ε* → 0 for bounded ranges of *K*.

## 5 Small-noise asymptotics

The master equation (7) can succinctly be written as a continuity equation

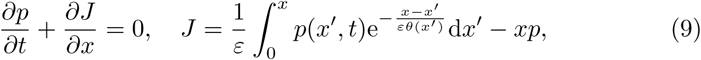

where

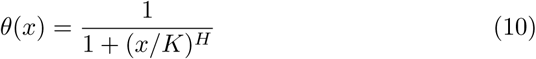
gives the factor by which the mean burst size is reduced at protein concentration *x* due to self-regulation. Being the product of the burst frequency *ε*^−1^and the mean burst size (6), it can also be interpreted as the mean protein production rate. The function *θ*(*x*) has a (unique) fixed point *x*_0_, i.e. a solution to

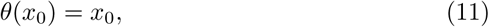
which can be interpreted as the protein concentration at which its mean production rate is equal to its rate of decay (cf. Fig. 1, right panel). It is shown below that in the small-noise regime the steady-state protein concentration is constrained with high probability to a small neighbourhood of the fixed point.

The stationary distribution is the time-independent, zero-flux, and normalised solution *p*(*x*) to (9), i.e the normalised solution to the integral equation

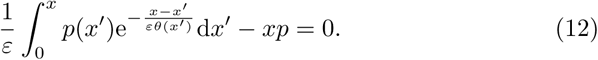

In order to examine the behaviour of the solution *p*(*x*) = *p*(*x*;*ε*) to (12) in the small-noise regime (*ε* ≪ 1), we use the transformation

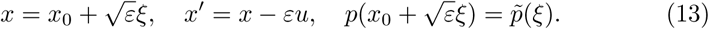

The substitution for *x* reflects the (presumed) 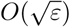 scale of fluctuations of *x* around the fixed point *x*_0_, cf. Eq. (5); the substitution for *x′* reflects the *O*(*ε*) scale of individual bursts. Inserting the substitutions (13) into the integral equation (12), we obtain

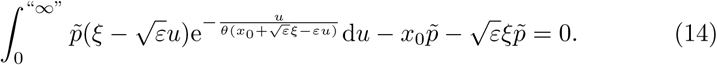

By extending the upper integration boundary from 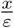 to infinity in (14), we will have introduced an error that is inconsequential (exponentially small) in the *ε* ≪ 1 regime (Hinch, 1991). Inserting the expansions

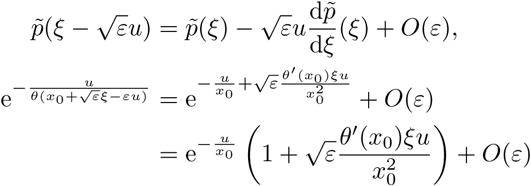
into (14) and carrying out the integration with respect to *u* we obtain

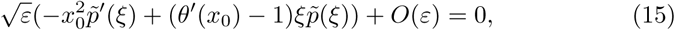
the *O*(1) terms having cancelled each other out. Dividing (15) by 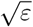 and neglecting the small 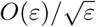 term, we arrive at a linear first-order differential equation for the leading-order approximation to 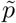, solving which yields

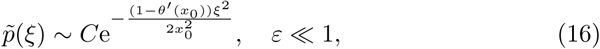
where *C* is the normalisation constant. The result implies that *ξ* is approximately normally distributed with mean zero and variance 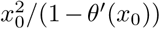. Consequently 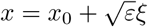 has mean *x*_0_ and variance 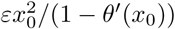, i.e.

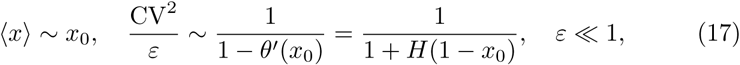
hold in the small-noise regime. Also, since the variance is *O*(*ε*), we obtain

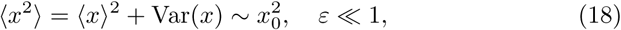
for the mean square.

In Fig. 5, we compare for *H* = 5 the small-noise approximations (17) (SNAs; dashed lines in Fig. 5) to the exact (i.e. not asymptotic) mean and relative CV^2^ for a range of values of *K* and three selected values of *ε*: a subcritical *ε* = 0.05, the critical *ε* = 0.1, and a supercritical *ε* = 0.2. The fixed point *x*_0_ of the function *θ*(*x*) (10) was calculated in the Python programming language with the `fixed_point` routine of the `scipy.optimize` library. For each value of *ε*, and for the range of values of *K*, the exact mean and the CV^2^ were estimated from the temporal average of a single and long sample path generated by Algorithm 1, which we implemented in the C programming language (see Appendix A for details).

**Figure 5:**
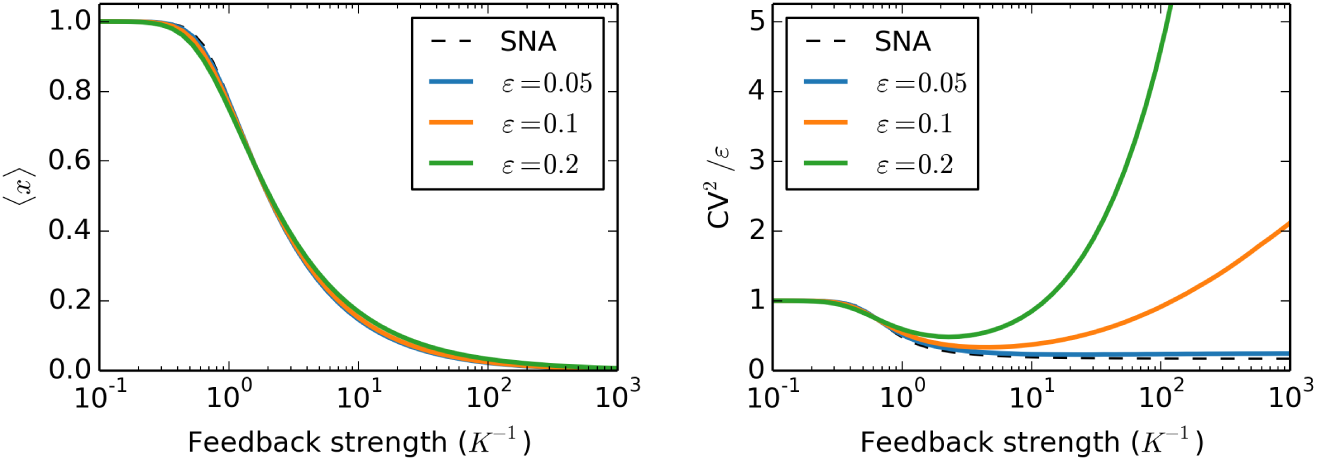
Protein concentration mean and the relative CV^2^in response to increasing feedback strength for *H* = 5 and a selection of values of *ε*, compared with the small-noise approximation (SNA).

By Fig. 5, left panel, the small-noise approximation of the mean agrees well with the exact mean, decreasing monotonically from one in the absence of regulation (*K* = ∞) to zero under stringent regulation (*K* → 0). Contrastingly, the small-noise approximation of the CV^2^ agrees with the exact CV^2^ only if *ε* is subcritical or *K* is sufficiently large; in particular, if *ε* is low and subcritical, the exact relative CV^2^ remains close, as *K* → 0, to the small-noise approximation limit value of 1/(*H* + 1) ≈ 0.167. For supercritical noise loads, however, simulations suggest divergence, as *K* tends to zero, of the exact CV^2^ to infinity, which the small-noise approximation fails to predict.

## 6 Strong-feedback asymptotics

In the small-noise analysis presented in the previous section, we tacitly assumed that the fixed point *x*_0_ of the Hill function (10) is of order one. However, *x*_0_ becomes increasingly small as *K* tends to zero; more precisely, the dominant balance in the fixed-point equation 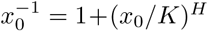 is found between the 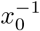 and the (*x_0_*/*K*)*^H^* terms, the equating of which yields a power-law relationship

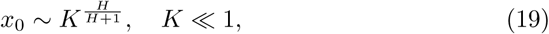
between the fixed-point *x*_0_ and the critical concentration *K* in the strong feedback regime. As *x*_0_ becomes increasingly small, the terms which have been neglected in the small-noise approximation procedure become significant, and the approximation leads to incorrect predictions; in particular, it fails to predict the grow-up of noise under supercritical noise-load conditions (Fig. 5, right panel).

In order to obtain a truthful characterisation of the model behaviour in the strong-feedback regime, we consider the steady-state protein probability density function in this section as a function of *K* which tends to zero, whereby the value of *ε* is fixed and not necessarily small. By (12) and (10), the steady-state protein pdf satisfies the integral equation

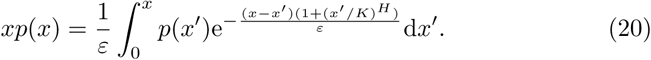

Below we show that a solution *p*(*x*) = *p*(*x;K*) to (20) tends to two separate distinguished limits, one occurring on the inner *x* = *O*(*K*^*H*/(*H* + 1)^) concentration scale associated with the fixed-point asymptotics (19), the other emerging on the outer *O*(1) scale associated with large, uncontrolled, bursts. The relative importance of the inner or outer limits will determine, in particular, whether the CV^2^ converges or diverges as *K* tends to zero.

### Inner solution

We rescale the variables of (20) according to

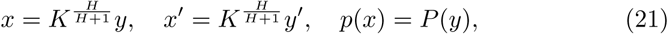
obtaining

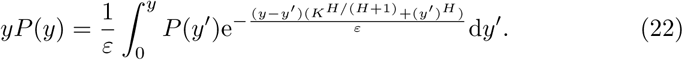

Inserting

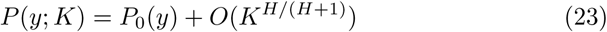
into (22) and collecting leading-order terms yield

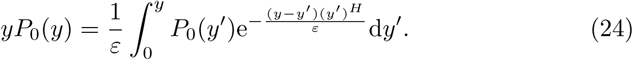

If *y* ≪ 1, then the exponential in the integrand is close to one, implying that *P*_0_(*y*) ∼ *ϕ*_0_(*y*), where *ϕ*_0_(*y*) satisfies 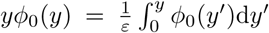, i.e. *ϕ*_0_(*y*) = *Cy*^1/*ε*−1^; for uniqueness and algebraic simplicity we fix the arbitrary constant *C* to one, thus obtaining

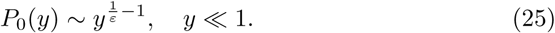

We refer to the (uniquely determined) solution to (24) which additionally satisfies (25) as the *inner solution*; it depends on *ε* and *H* but is independent of the value of *K*.

The integral equation (20) determines an exact solution *p*(*x*) up to a multiplicative constant. The inner solution approximates the exact solution which additionally satisfies

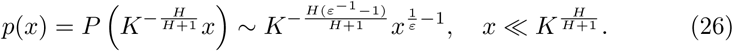

The (uniquely determined) solution *p*(*x*) to (20) and (26) is not a probability density function, as it does not necessarily integrate to one. Prior to normalising the solution, we characterise its approximation on the outer scale and evaluate the asymptotic behaviour of its moments.

### Outer solution

We substitute *x′* = *Kz* into (20), while leaving *x* = *O*(1), which gives

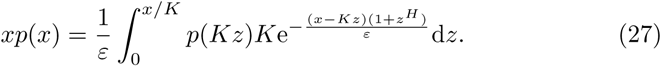

Recalling that *x′* has the interpretation of the protein concentration immediately before a burst and *x* represents (a lower bound for) the concentration after the burst, the current scaling relates to the possibility that the protein may deviate to very low *O*(*K*) levels, at which the burst size ceases to be under regulation, hence the following burst is *O*(1) large.

Since the *O*(*K*) concentration scale is smaller than the *O*(*K*^*H*/(*H*+1)^) inner concentration scale, we can approximate *p*(*Kz*) on the right-hand side of (27) by the asymptotic expansion (26), i.e.

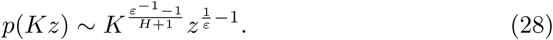

We insert (28) into (27), extend the upper limit of integration in (27) to infinity and neglect small *O*(*K*) terms in the argument of the exponential in (27), obtaining

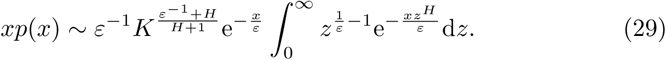

Substituting *u* = *xz^H^*/*ε* in the integral in (29) and dividing both sides of the relation by *x*, we obtain

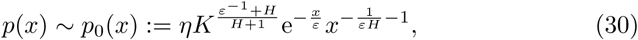
where

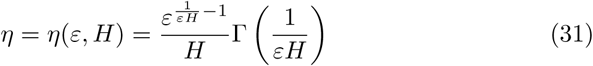
is a constant which depends on *ε* and *H* only. We refer to the function *p*_0_(*x*) defined by the right-hand side of (30) as the *outer solution*.

### Approximating the moments

The *n*-th moment of the solution *p*(*x*) to (20) satisfying (26) is defined by

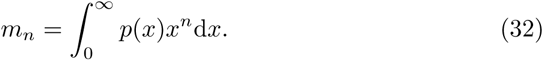

By (30), the outer approximation *p*_0_(*x*)*x^n^* of the integrand in (32) is of the order of 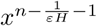, as *x* tends to zero; if 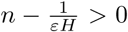, it is integrable: it increases too slowly as *x* → 0 for the boundary layer to make a significant contribution to the integral (Hinch, 1991). Thus, the *n*-th moment, for 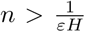, of the exact solution *p*(*x*) can be approximated for small values of *K* by the *n*-th moment of its outer approximation (30), i.e.

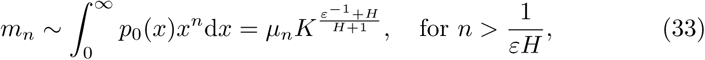
where

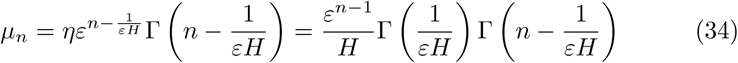
is a constant which does not depend on *K*.

Substituting 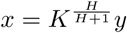 in (32), we express *m_n_* for 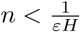 in terms of the inner variable,

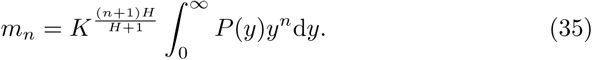

By asymptotic matching (Kevorkian and Cole, 1981), the asymptotic behaviour of the inner solution *P*_0_(*y*) as *y* → ∞ coincides with that of the outer solution *p*_0_(*x*) as *x* → 0. Consequently, the inner approximation *p*_0_(*y*)*y^n^* of the integrand in (35) is of the order of 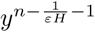 as *y* → ∞. If 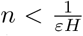, the inner solution is integrable, decreasing too rapidly as *y* → ∞. for the outer region to contribute significantly to the integral (Hinch, 1991).

Thus, we can write

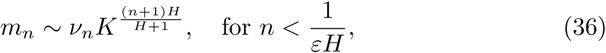
where

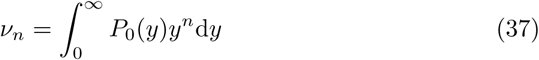
is the *n*-th moment of the inner solution, which does not depend on *K* and depends on *ε* and *H* only.

### Normalisation

The normalised steady-state protein pdf is obtained by dividing the solution *p*(*x*) to (20) satisfying (26) by its norm 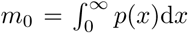. Combining (26), (30) and (36) we arrive at asymptotic approximations

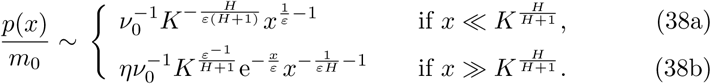

Figure 6 shows an excellent agreement between the theoretical scalings (38a)–(38b) and probability distributions estimated by extensive kinetic Monte Carlo simulation with Algorithm 1.

**Figure 6:**
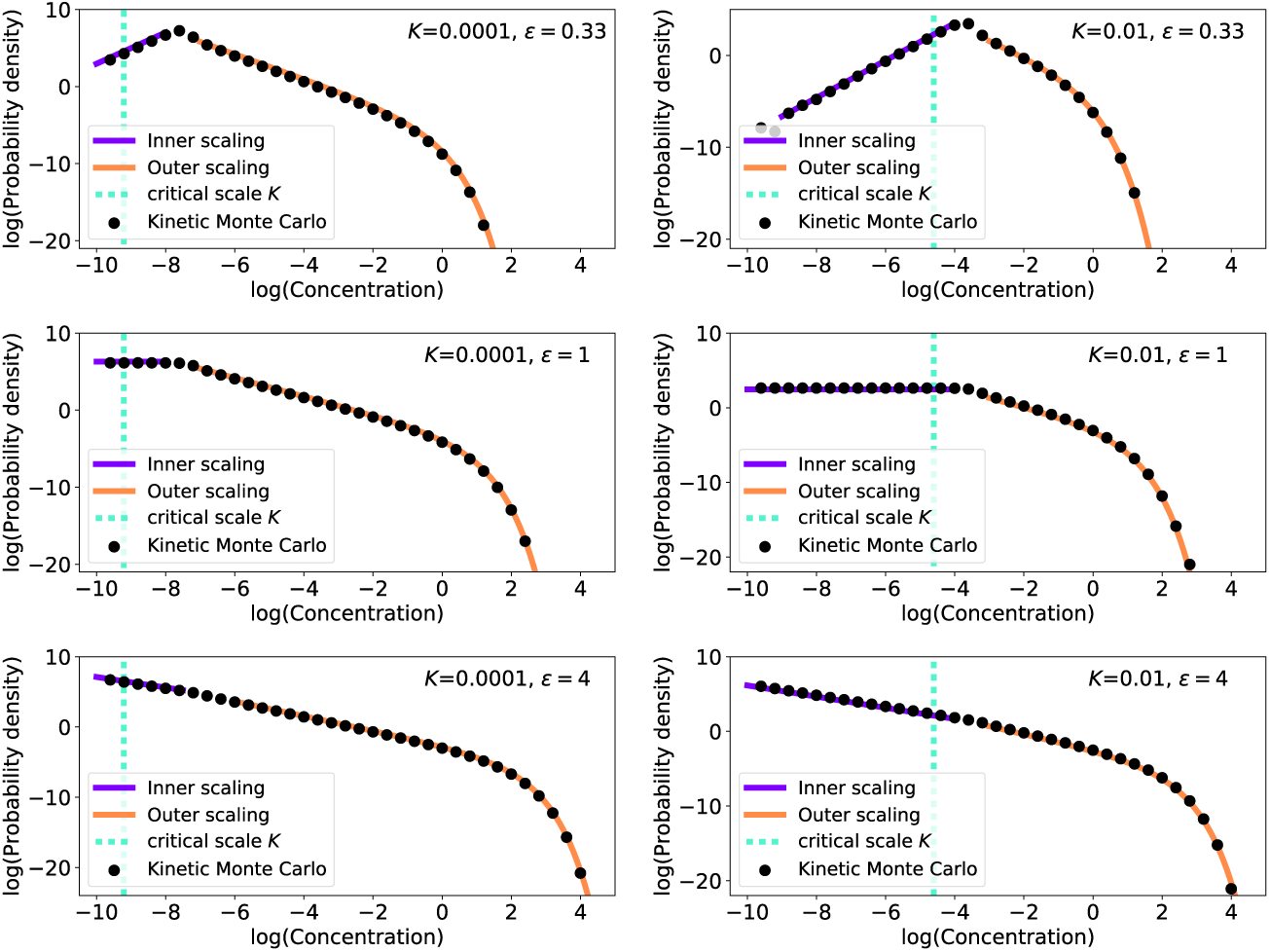
Protein distributions for selected parameter sets. Discrete markers: probability distributions estimated by kinetic Monte Carlo simulation with Algorithm 1; dashed line: the critical concentration *K*; solid lines: theoretical predictions for the inner (38a) (purple colour) and outer (38b) (orange colour) scalings of the probability distributions with manually tuned prefactors 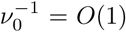 The simulation data were estimated from 10^6^ sample paths after the stationarities of the process were reached.

### Approximating the protein mean and CV^2^

Using the asymptotic resuits (33) and (36) for the moments *m*_0_, *m*_1_, and *m*_2_, we can also derive approximations for the mean and the mean square, which are given by the ratios

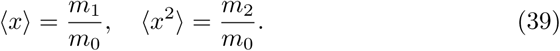

The mean satisfies, as *K* → 0,

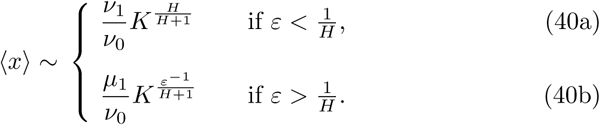

By (40a), under low-noise conditions 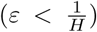, the mean exhibits the same power-law decrease as *K* → 0 as its small-noise prediction *x*_0_ (19), albeit with a prefactor which is different from one. As *ε* exceeds the threshold 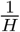, the exponent of the power law (40b) becomes smaller with increasing *ε*, implying a slower decrease of the mean as *K* → 0 as that predicted by the small-noise approximation.

For the mean square we obtain in the small-*K* regime the asymptotics

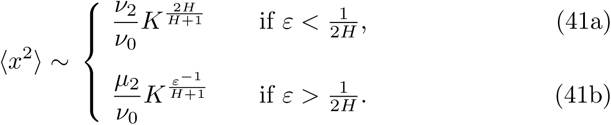

Thus, for small values of *ε*, the mean square 〈*x*^2^〉 decreases as *K* → 0 with the same exponent 2*H*/(*H* + 1) as its small-noise prediction 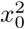 (18). The alternative, slower, exponent *ε*^−1^/(*H* + 1) applies if *ε* exceeds the threshold 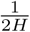, which is half the threshold in (41) for the mean and which we previously in Section 4 referred to as the critical noise load.

The coefficient of variation can be approximated for small values of *K* by

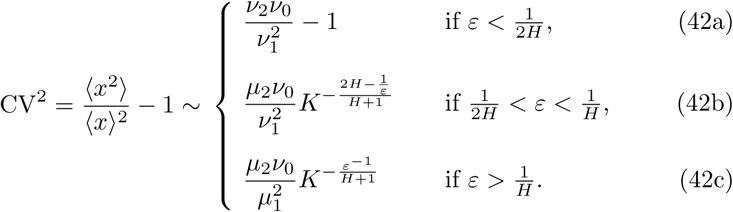

By (42a), CV^2^ converges to a constant value as *K* → 0 for subcritical noise loads 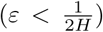. For supercritical noise loads 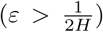, the CV^2^ exhibits a power-law increase as *K* decreases to zero. As *ε* increases beyond the critical value 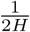, the absolute value of the exponent of the power law in (42b) first increases, reaching a maximum of *H*/(*H* + 1) for 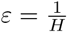; after *ε* exceeds 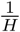, the exponent of the power law in (42c) begins to decrease in absolute value. Thus, the fastest grow-up in the coefficient of variation as *K* → 0 is achieved for 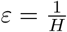 by a combination of a relatively fast decrease in the mean and a relatively slow decrease in the variance. Under excessive noise conditions 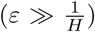, feedback plays a limited role, which is evidenced by low grow-up rates of CV^2^.

For numerical analysis, we fixed *H* = 5, used a selection of 20 values for *ε* ranging from 0.05 to 1, and a geometric sequence of 5 values for *K* ranging from 10^−7^ to 10^−3^. For each of the 20 × 5 parameter combinations from the (*H, ε, K*) parameter space of the model, we estimated the steady-state mean concentration value 〈*x*〉 and the mean square value 〈*x^2^*〉 by generating 10^6^ sample paths using Algorithm 1. The *ensemble average* of the first and the second moments were measured after stationarity of the process is reached, after which time the moments were measured every 0.2 units of time for another 5 x 10^3^ units of time to perform an additional *temporal averaging*.

The simulation results are visualised in Fig. 7, where we show the squared mean 〈*x*〉^2^ (Fig. 7, top left), the mean square 〈*x*^2^〉 (Fig. 7, top right), and the CV^2^ (Fig. 7, centre left) as functions of the critical concentration *K*. We use (decimal) logarithmic scale for both axes in all panels; any power-law relationship then appears as a straight line with slope which is equal to the power-law’s exponent. Using simple linear regression, we estimate the power-law exponents (Fig. 7, bottom, shown as triangular and square markers) and compare the estimates to the asymptotic predictions (40)–(42) (Fig. 7, bottom, black lines).

**Figure 7:**
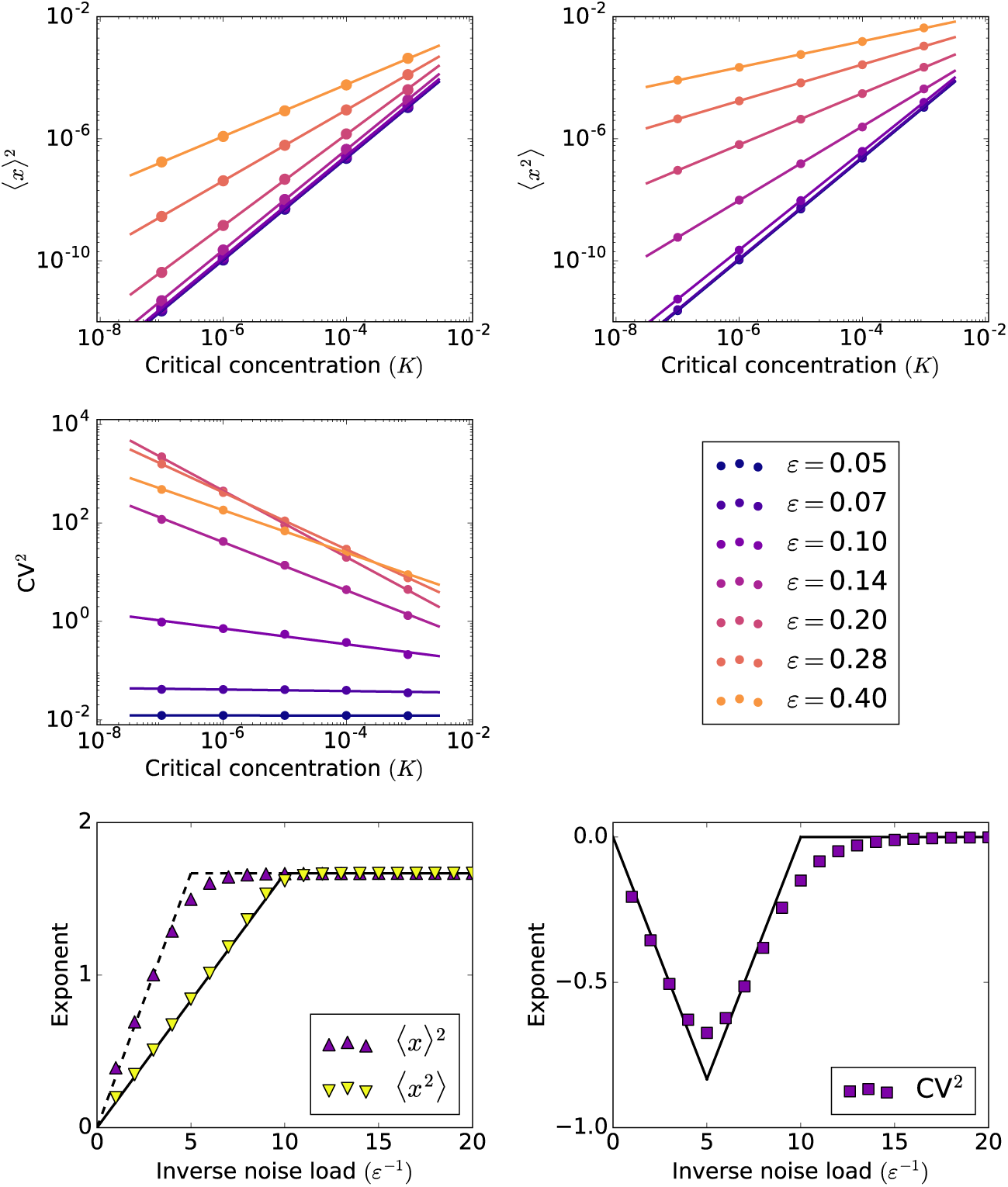
The squared mean (top left), the mean square (top right) and the CV^2^ (centre left), obtained by stochastic simulation (Algorithm 1), as functions of critical concentrations *K* ≪ 1 for *H* = 5 and a selection of values of *ε*(centre right). The discrete markers are the numerically measured results. The exponents of the power-law dependence on *K* of 〈*x*〉^2^, 〈*x^2^*〉 (bottom left), and CV^2^ (bottom right), obtained by linear regression (solid lines in top left, top right, and centre left) and asymptotic analysis (40)–(42), as functions of the inverse noise load *ε*^−1^.

While the simulation-based exponent estimates in Fig. 7 are in a general agreement with the asymptotic results for 〈*x*〉 in (40) and 〈*x*^2^〉 in (41), some discrepancies are observed around the critical values of *ε*(which are *ε* = 0.2 for 〈*x*〉^2^ and *ε* = 0.1 for 〈*x*^2^〉). We attribute these errors to logarithmic correction terms which appear in the leading-order asymptotic approximations for the moments at critical values (see Appendix B).

The exponent for the CV^2^ is given by the difference of the exponents for 〈*x*^2^〉 and that for 〈*x*〉^2^. The simulation-based estimate of the exponent for the CV^2^ incorporates the errors incurred in estimating the exponents for both 〈*x*^2^〉 and that for 〈*x*〉^2^ : the resulting error can be quite large relative to the absolute value of the exponent itself. Additionally, the constant term −1, which we neglected as higher-order in the asymptotic approximations (42b) and (42c), can nevertheless be relatively important if the absolute value of the exponent is low; this adds yet another source of error in the estimate of the exponent. Despite these inaccuracies, the simulation-based results for the CV^2^ are in qualitative agreement with the leading-order asymptotic predictions: the CV^2^ remains bounded if *ε* < 0.1 and exhibits the fastest grow-up for *ε* = 0.2.

## 7 Comparison with previous results

In our present model for feedback in burst size, the concentration of protein immediately before a burst occurs determines the expected size of the burst. The distribution of burst sizes is exponential, whose defining property is its memorylessness: at any stage of the growth of a burst, the amount of protein yet to be produced is independent of how much has already been produced. Such lack of self-control on a single-burst level implies that our model implicitly includes a delaying step, which newly produced molecules have to undergo before they can take part in the self-regulation. In order that our model be applicable, the delay must be neither too short nor too long: on one hand, it needs to last long enough to carry over the duration of a single burst; on the other hand, it must be short enough so that all protein produced in the current burst will have maturated by the time of the next burst. Biologically, such delays can easily be accounted by the time it takes to complete the synthesis of a gene product. As a specific example, translational bursts in prokaryotes occur when a short-lived mRNA is transcribed from a gene and repeatedly translated before it is degraded (McAdams and Arkin, 1997). Each translation begins when the mRNA is bound by a ribosome and continues by the ribosome sliding down the mRNA molecule. As it moves along the mRNA, the ribosome forms an elongating chain of amino acids, which is to become a protein after the ribosome reaches the end of the mRNA code. Since the binding of new ribosome molecules occurs while the previous ones are still elongating, the proteins whose translation initiated earlier cannot inhibit those initiated later within a single burst.

In an alternative version of our model for feedback in burst size, which was analysed in a previous paper (Bokes and Singh, 2016), the growth of a burst depends on the current value of the protein concentration, including the molecules already synthesised within the same burst, rather than on its pre-burst level. Consequently, the distribution of burst sizes is not exponential and harder to draw random variates from, which complicates the implementation of a fast and exact simulation algorithm. On the other hand, the model in (Bokes and Singh, 2016) is considerably easier to treat analytically: the master equation of the undelayed model reads

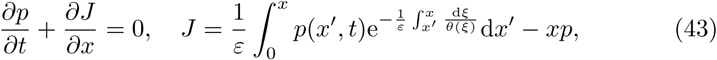
where *θ*(*x*) = (1 + (*x*/*K*)^*H*^)^−1^ is the Hill function. The integral term in the probability flux *J* can be recognised as the variation-of-constants solution *f*(*x*) to the differential equation *df*/*dx* + *f*/*εθ* = *p* subject to *f*(0) = 0. Applying the differential operator *d*/*dx* + 1/*εθ* on the steady state equation *J* = 0 transforms this integral equation into an ordinary differential equation (Lin and Doering, 2016), solving which yields an explicit steady-state probability density function

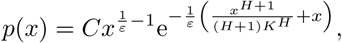
where *C* is a normalisation constant. Steady-state protein moments can be evaluated by numerical integration of the density or by asymptotic approximation of the integrals in the small-noise and strong feedback regimes (Bokes and Singh, 2016). In the small-noise limit, the results that follow are identical with (17) obtained for the current model (Fig. 8, left and central panels, dashed lines). The non-delayed model exhibits a monotonic decrease in the coefficient of variation for any combination of *ε* and *H* (Fig. 8, central panel, coloured lines). The loss of control over protein noise, which occurs for supercritical noise loads in the strong-feedback regime of our current model (Fig. 8, left panel, coloured lines), can therefore be attributed to the delay which carries over the bursting timescale.

**Figure 8:**
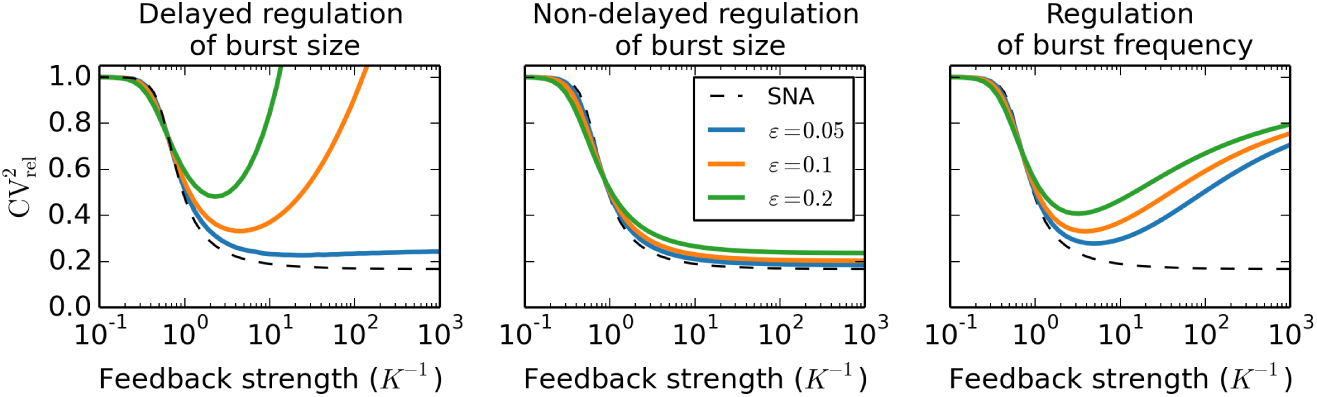
The relative CV^2^ of a protein produced in bursts subject to different types of feedback in response to increasing feedback strength *K*^−1^. We fix *H* = 5 and use a selection of values of *ε* which are given in the legend within the central panel. Exact numerical/simulational results are compared to the small-noise approximation (SNA).

While so far we have focused exclusively on feedback in burst size, more attention has traditionally been paid in literature to feedback in burst frequency. If one adheres to the fundamentals of the model of Section 2, but uses the Hill function to reduce the burst frequency in the excess of protein, rather than reducing the expected burst size, one arrives at a model for negative feedback in burst frequency which was also analysed in (Bokes and Singh, 2016). The master equation of the model reads

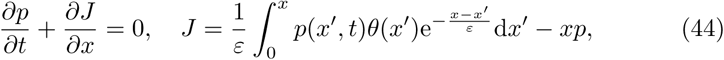
which admits an explicit steady-state solution, see (Lin and Doering, 2016; Bokes and Singh, 2016; Friedman et al, 2006), given by

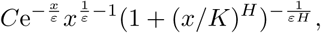
where *C* is a normalisation constant. In the small-noise limit, the steady-state mean converges to the fixed point *x*_0_ of the Hill function *θ*(*x*) = (1+(*x*/*K*)^*H*^)^−1^ consult Bokes and Singh (2016) for details, which is the same value as was obtained in (17) for feedback in burst size. The squared coefficient of variation, on the other hand, was found in (Bokes and Singh, 2016) to be larger by a factor of 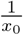 than the value (17) obtained for feedback in burst size. We should note, however, that, even in the absence of any regulation, the squared coefficient of variation is inversely proportional to the burst frequency (cf. Section 2). The increase in the coefficient of variation in the small-noise limit can therefore be attributed to the a decrease in the overall (time-averaged) burst frequency from the unregulated value of 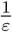 to the regulated value of 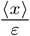 rather than to a loss of control over protein noise by the regulation of burst frequency.

Adjusting for the decrease in the overall burst frequency, Bokes and Singh (2016) defined the relative coefficient of variation as the ratio of the coefficient of variation of the self-regulating protein and that of a constitutively expressed protein with the same overall, time-averaged, frequency of bursts. While for the feedback in burst size this definition trivially reduces to 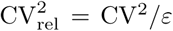, for feedback in burst frequency, the relative coefficient of variation is given by 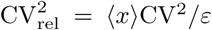. In the small-noise limit, both feedback types yield the same approximation for the relative coefficient of variation, which decreases monotonically with increasing feedback strength (Fig. 8, dashed lines). For feedback in burst frequency, the relative coefficient of variation diverges for *K* ≪ 1 from the small-noise prediction, but in a manner which is qualitatively different from the one which we identified in this paper to occur for feedback in burst size (Fig. 8, left and right panels, coloured lines).

Consistently with the small-noise prediction, the relative coefficient of variation initially decreases from the value of one as feedback in burst frequency strengthens, until the critical concentration *K* drops to *O*(*ε*) levels, at which point the relative coefficient of variation begins to increase again, converging back to one as *K* tends to zero (Fig. 8, right panel; consult Bokes and Singh (2016) for asymptotics). Although at lower noise loads the turnaround occurs at higher feedback strengths, a critical noise load below which the small-noise approximation would be uniform does not exist, and an eventual loss of control over noise is inevitable in the small-*K* limit (Fig. 8, right panel). On the other hand, a protein which regulates its burst frequency is never noisier than a constitutively expressed with the same average burst frequency, whereas a protein with feedback in burst size will indeed be noisier if subjected to supercritical noise loads. Bokes and Singh (2016) previously compared the-noise reduction performance of feedback in burst frequency and (undelayed) feedback in burst size, reporting that the latter always performs better. In light of the present results, we conclude that depending on parametric conditions such as indicated above, either (delayed) feedback in burst size or in burst frequency can be optimal in terms of minimising the noise.

## 8 Discussion

The synthesis of protein molecules has been shown to occur in bursts of rapid production which alternate with periods of inactivity. In a minimalistic model for burst-like protein expression, bursts are represented by randomly occurring, randomly sized, discontinuous jumps in protein concentration; these are counterbalanced by deterministic decay of the concentration due dilution by cell growth and/or active degradation. We extended this minimalistic model by a specific kind of negative feedback, making the expected burst size decrease with increasing protein concentration. We investigated the ability of such kind of negative feedback to control stochastic variability in protein levels.

Three positive dimensionless parameters — the noise load *ε*, the cooperativity coefficient *H*, and the critical protein concentration *K* — completely determine the behaviour of the model at steady state. The noise load *ε* is equal to the squared coefficient of variation the protein exhibits without any feedback. Its reciprocal *ε*^−1^ is equal to the average number of bursts per protein lifetime. The cooperativity coefficient *H* measures the steepness of decrease in the expected burst size in response to increasing protein concentration. Biologically, a positive integer value of *H* means that *H* protein molecules interact to form a complex which interferes with the transcription or translation machinery to reduce the burst size. The critical concentration *K*, measured in the units of the protein concentration mean in the absence of regulation, gives the threshold value which is required to reduce the expected burst size by half. We use the parameter *K* as an inverse measure of feedback strength: the lower the threshold for efficient self-repression, the stronger the feedback.

The central result of the paper lies in characterising the response of the steady-state protein statistics — the coefficient of variation in particular — to the strengthening of feedback, i.e. to decreasing *K*, while keeping the noise load *ε* and the cooperativity coefficient *H* constant. We found dramatically different responses depending on whether the noise load is less than or greater than a critical value 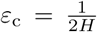. For subcritical noise loads, the coefficient of variation remains bounded with increasing feedback strength. Contrastingly, for supercritical noise loads, the coefficient of variation diverges to infinity as a power of *K* with a negative exponent.

The identification of the critical noise load is one important consequence of a bi-scale behaviour of the protein probability density function *p*(*x*) = *p*(*x*; *K*) as *K* → 0 characterised in Section 6. In addition to the outer *x* = *O*(1) scale of large uncontrolled bursts, there is also an inner *x* = *O*(*K*^*H*/(1+*H*)^) scale of small controlled bursts, on which the density tends to two distinguished limits, the outer and the inner solutions. The small-*K* behaviour of the moments 〈*x^n^*〉 is decided in the overlap region *K*^*H*/(1+*H*)^ ≪ *x* ≪ 1, in which the density is proportional to the power 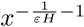. The integrability of 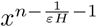. as *x* → 0 (if 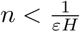) or as *x* → 0 (if 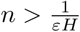) decides whether the dominant contribution to the *n*-th moment comes from the inner or the outer region. If the inner region is dominant, the *n*-th moment tends to zero as *K*^*nH*/(1+*H*)^; if the outer region dominates, the convergence of the *n*-th moment is still polynomial but slower. The critical exponent 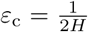 thus gives the value of noise value at which the second moment ceases to be determined in the inner region of controlled bursts and begins to be driven by the outer region of uncontrolled bursts. The small-*K* behaviour of the second moment, and to a lesser extend that of the first moment (the mean), determine whether the coefficient of variation converges or diverges, as well as the rate of the divergence, as *K* tends to zero. We anticipate a potential criticism that our emphasis on the coefficient of variation is arbitrary and obscures the effect of the noise load and cooperativity coefficient on the polynomial decay of the density in the overlap region. Nevertheless, the coefficient of variation has widely been used both in experimental and theoretical analyses of stochastic gene expression. By focusing on the well-known measure of noise, we make our asymptotic analysis relatable to previous work and potentially interesting to a wider audience.

Our results have implications for the role of high cooperativity in negative feedback systems. Using a standard small-noise (*ε* ≪ 1) approximation approach, we have shown that feedback in burst size can reduce the squared coefficient of variation by a factor of *H* + 1, thus confirming previous reports that cooperativity leads to improved attenuation of noise (Singh and Hespanha, 2009). This observation is not specific to feedback in burst size but also holds for feedback in burst frequency (Bokes and Singh, 2016) provided that one compensates for the drop in the overall burst frequency, which is larger for cooperative feedbacks. Contrastingly, using an alternative strong-feedback (*K* ≪ 1) approximation approach, we have identified, specifically for feedback in burst size, an adverse effect of cooperativity on protein noise: high cooperativity can lead to a significant amplification of protein noise in response to increasing feedback strength even if the underlying noise load is relatively low 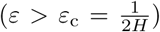. It is well known that high-cooperativity feedbacks are prone to instability if they operate with a sufficiently large delay (Murray, 2003). Interestingly, the loss of control over noise in our present model can also be attributed to a delay, albeit an infinitesimally small one: it has been introduced into the model by assuming that the mean burst size is determined by the protein concentration immediately before the burst starts. We expect that the interesting interplay between bursting noise, cooperativity, and delay, which we have illustrated here within a minimalistic modelling framework, will have implications in other, more complex, systems also.

## Appendix A. Estimating protein moments from simulations

We can estimate the *n*-th steady-state moment by the time average 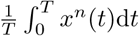, where *T* ≫ 1, of a sample trajectory *x*(*t*) generated by Algorithm 1 and raised to the power of *n*. Since *x*(*t*) is piecewise exponential, we have

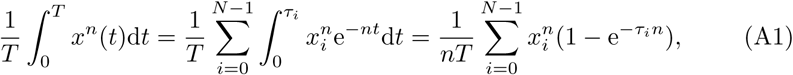
where *x_i_, i* ≥ 1, is the protein concentration immediately after the *i*-th burst, *x*_0_ is the initial protein concentration, *τ_i_, i* ≤ 1, is the waiting time from the *i*-th burst until the (*i* + 1)-th burst, *τ*_0_ is the waiting time from the initial time until the first burst, and 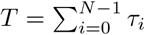, where *N* is a large integer. By Algorithm 1, the values of *x_i_* and *τ_i_* are obtained by

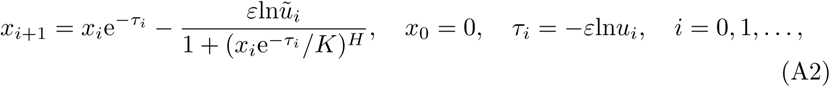
where *u_i_* and *ũ_i_, i* = 0,1,…, are random variates drawn independently of each other from the uniform distribution in the unit interval.

Inserting *T* ≈ *εN*, which holds by the law of large numbers, into (A1), and shifting the time frame to reduce the effect of the transient behaviour, we arrive at an estimate

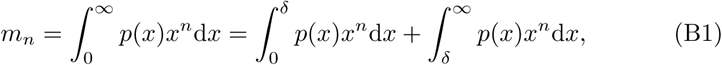
where *x_i_* and *τ_i_* are given by (A2). We used (A3) with *N* = 10^9^ and *M* = 10^7^ to estimate the theoretical value of 〈*x^n^*〉 in Figures 3–5 and Fig. 8. In the strong feedback limit (Figures 6 and 7), ensemble averaging across a large number of sample paths was needed for more precise measurements of the power-law exponents.

## Appendix B. Strong-feedback asymptotics in critical cases

The dominant contribution to the *n*-th moment of a solution *p*(*x*) to (20) comes exclusively from the inner *O*(*K*^*H*/(*H*+1))^ concentration scale if 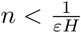 or from the outer *O*(1) concentration scale if 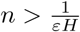. In the borderline case of 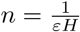, which we explore in this Appendix, the inner and outer regions both contribute to the leading-order behaviour of the *n*-th moment. The individual contributions can be identified by splitting the range of integration (Hinch, 1991),

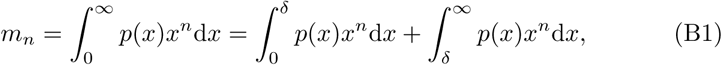
where *δ* is a value taken from an intermediate scale (*K*^*H*/(*H*+1)^ ≪ *δ* ≪ 1).

In the first integral on the right-hand side of (B1), we substitute *x* = *K*^*H*/(*H*+1)^*y* and replace the integrand by the inner approximation,

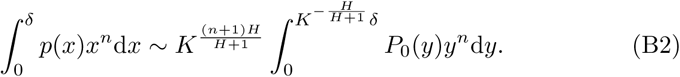

Since *δ* ≪ *K*^*H*/(*H*+1)^, the upper integration limit of the integral on the right-hand side of (B2) tends to infinity as *K* becomes increasingly small. The integral itself then diverges to infinity, since by (30)–(31) and van Dyke’s matching rule an asymptotic expansion

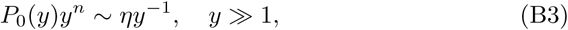
holds for the integrand. Extricating the divergent part from the integral in (B2), we obtain an asymptotic approximation

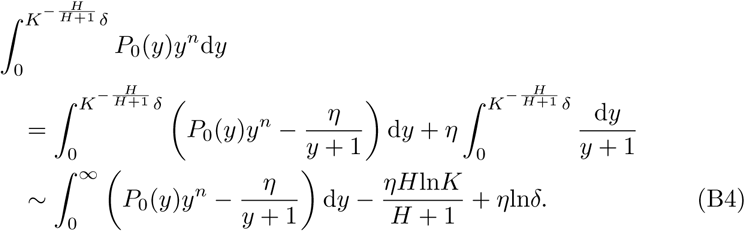

In the second integral on the right-hand side of (B1), we replace the integrand by the outer approximation (30), obtaining

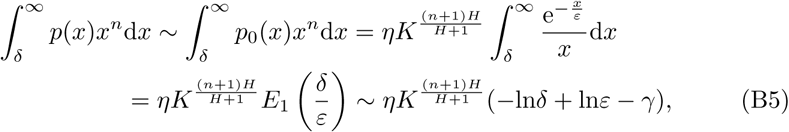
where *E*_1_(*t*) is the exponential integral and *γ* = 0.577 … is the Euler–Mascheroni constant.

Collecting (B2), (B4), and (B5), we find that the *n*-the moment can be expanded into

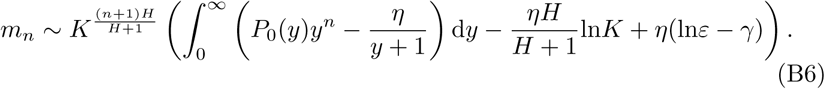

Although the logarithmic term in (B6) asymptotically dominates, as *K* tends to zero, the neighbouring constant terms, in practice the magnitudes of the logarithm and the constant terms are similar so the latter cannot be neglected.

As a particular application of the expansion (B6), we evaluate the small-*K* behaviour of the protein coefficient of variation subject to a critical noise load *ε* = 1/2*H*. The leading-order approximations to *m*_0_ and *m*_1_ involve contributions from the inner scale only and are given by (36)–(37). On the other hand, the leading-order approximation to *m*_2_ combines contributions from either scale and is given by (B6). For the coefficient of variation we obtain

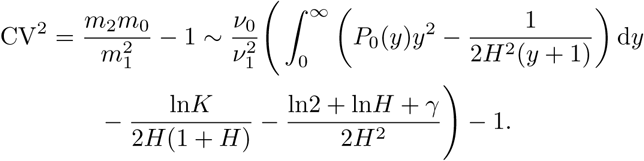

In the Main Text, we showed that the coefficient of variation remains bounded as *K* goes to zero if *ε* < 1/2*H* and increases polynomially if *ε* > 1/2*H*. The above result implies that in the critical case *ε* = 1/2*H* exhibits a slow logarithmic increase as *K* tends to zero.

